# Determination of phylogenetic relationships in the genus *Mangifera* based on whole chloroplast genome and nuclear genome sequences

**DOI:** 10.1101/2023.09.24.559211

**Authors:** WWM Upendra Kumari Wijesundara, Agnelo Furtado, Natalie L. Dillon, Ardashir Kharabian Masouleh, Robert J Henry

**Author notes:** **Corresponding Author**: Robert J. Henry.

## Abstract

The genus *Mangifera* (*Anacardiaceae*) includes 69 species of which *Mangifera indica L*. is the most important and primarily cultivated species for commercial mango production. Although the species are classified based on morphological descriptors, molecular evidence has proposed the hybrid origin of two species suggesting the possibility that more of the species may be of hybrid origin. To analyze evolutionary relationships within the genus, 13 samples representing 11 *Mangifera* species were sequenced and whole chloroplast (Cp) genomes and 47 common single-copy nuclear gene sequences were assembled and used for phylogenetic analysis using concatenation and coalescence-based methods. The Cp genome size varied from 151,752 to 158,965 bp with *M. caesia* and *M. laurina* having the smallest and largest genomes, respectively. Genome annotation revealed 80 protein-coding genes, 31 tRNA and four rRNA genes across all the species. Comparative analysis of whole Cp genome sequence and nuclear gene-based phylogenies revealed topological conflicts suggesting chloroplast capture or cross hybridization. The Cp genomes of *M. altissima*, *M. applanata*, *M. caloneura* and *M. lalijiwa* were similar to those of *M. indica* (99.9% sequence similarity). Their close sequence relationship suggests a common ancestry and likely cross-hybridization between wild relatives and *M. indica*. This study provides improved knowledge of phylogenetic relationships in *Mangifera,* indicating extensive gene flow among the different species, suggesting that hybrids may be common within the genu*s*.

## 1. Introduction

Mango (*Mangifera indica* L), an evergreen dicotyledonous angiosperm often referred as “king of fruits” is adapted to grow in tropical and sub-tropical regions of the world (Mukherjee 1949a; Rabah et al. 2017; Singh et al. 2016; Vasanthaiah et al. 2007). It is considered as one of the most economically successful fresh fruits cultivated in more than 100 countries, with 55 million tonnes produced in 2020. India has produced approximately 24.7 million tonnes as the world’s largest mango producer accounting for 45% total mango production followed by Indonesia (6.6%), Mexico (4.3%), China (4.3%), and Pakistan (4.3%) (FAOSTAT 2022). Besides being consumed fresh, ripe and unripe mangoes are used to produce pickles, chutney, juices, cereal flakes, sauce, and jam building high demand for mangoes on the international market (Saúco 2016).

The taxonomic history of the genus *Mangifera* (*Anacardiaceae*) reveals consistent recognition of two major groups with the number of species reported varying between 45 -69 (Hou 1978; Kostermans and Bompard 1993; Mukherjee 1949a). The most accepted classification described by (Kostermans and Bompard 1993) defines 69 species mainly based on morphological descriptors of reproductive tissues. Of the 69 species, 58 are divided into two subgenera, *Mangifera* and *Limus*, with the remaining 11 species placed in an uncertain position due to insufficient voucher material. The subgenus *Mangifera* is characterized by free stamen filaments and a four/five lobed cushion-shaped papillose disc broader than the base of the ovary while and the subgenus *Limus* is characterized by united stamen filaments and a stalk-like disc narrower than the base of the ovary. The sub genus *Mangifera* includes 47 species further divided into four sections: *Marchandora* Pierre, *Euantherae* Pierre, *Rawa* Kosterm, and *Mangifera* Ding. In the subgenus *Mangifera*, section *Marchandora* Pierre includes only *M. gedebe* which has the unique character of labyrinthine seed. The three species (*M. pentandra* Hooker, *M. cochinchinensis* Engler, *M. caloneura* Kurz) belonging to section *Euantherae* Pierre are characterized by five fertile stamens. The section *Rawa* Kosterm includes nine species which are not well delimited. *Mangifera* Ding Hou is the largest section in the genus with more than 30 species including domesticated mango (*M. indica*) (Bompard 2009; Kostermans and Bompard 1993). The 11 species in sub genus *Limus* are further divided into two sections: *Deciduae* (deciduous trees) and *Perrennis* (non-deciduous species).

Due to the high demand for mango globally, systemic breeding programs have been initiated recently to develop cultivars with high productivity, consumer, and transportability traits more suited for national and international markets. However, breeding is time-consuming due to the long juvenile period, high heterozygosity and polyembryony observed in mango (Bally and Dillon 2018). Currently, although *M. indica* is the principal cultivated species for commercial fruit production (Dinesh et al. 2011) from which a set of selected commercial cultivars dominate the crop improvement programs, 26 other species have been reported to produce edible fruits including *M. altissima*, *M. foetida*, *M*. *caesia*, *M*. *odorata*, *M*. *pentandra*, *M*. *laurina, M. sylvatica, M. zeylanica* and *M*. *pajang* (Bally et al. 2021; Bompard 2009; Mukherjee and Litz 2009). Many wild species exhibit potential significance in trait-specific breeding due to their favourable traits related to fruit quality, biotic and abiotic stress tolerance and potential as rootstocks (Bompard 1992; Eiadthong et al. 1999; Iyer 1989) if their distinctive characteristics are properly exploited. Although the species have been described in terms of morphological characteristics, they are not well-characterized in a genetic framework. Therefore, identification of molecular evolutionary relationships within the genus is vital to allow efficient use of wild relatives in future breeding programs.

Recent studies have used molecular markers targeting coding and non-coding regions in the chloroplast genome (Eiadthong et al. 1999; Fitmawati et al. 2017; Hartana 2010; Hidayat et al. 2011) and a set of nuclear genes (Fitmawati 2016; Schnell and Knight Jr 1992; Yonemori et al. 2002) to analyse phylogenetic relationships within the genus. However, the results have not been consistent, and many studies were unsuccessful in inferring evolutionary relationships with fully resolved phylogenies. This may be due to use of molecular markers with limited genetic information within the genome, a slow evolutionary rate within the targeted regions and use of different taxa in different analysis. Two studies have used whole chloroplast genome (Niu et al. 2021) and mitochondrial genome (Niu et al. 2022) sequences alone with a small number of taxa. However, in most angiosperms, chloroplast and mitochondrial genomes are maternally and paternally inherited, respectively (Corriveau and Coleman 1988). These studies prevent precise analysis of evolutionary relationships due to the use of uniparentally inherited genetic information for phylogenetic analysis.

The genus *Mangifera* is native to South and South-East Asia ranging from Indochina, Burma, Thailand and the Malay Peninsula to Indonesia and Philippines where some of the species are found only in the wild while others are locally grown in gardens and orchards (Kostermans and Bompard 1993). With the introduction of common mango to South-East Asia during 4^th^-5^th^ century (Mukherjee 1949b), *M. indica* and wild *Mangifera* species in the region might have come into contact with each other. Since both wild and domesticated mangoes are assumed to be self-incompatible, hybridization is expected among these outcrossing species when grown in close proximity. Among wild species, a hybrid origin has been reported for *M. odorata* (Teo et al. 2002) and *M. casturi* (Matra et al. 2021; Warschefsky 2018). With molecular data suggesting the potential of cross-hybridization in the genus, more hybrids can be expected among these 69 *Mangifera* species that have been currently identified as distinct species. Comparative phylogenetic analysis based on both chloroplast genome and ideally, a set of single-copy nuclear genes, together representing maternal and biparental inheritance respectively (Liu et al. 2020; Tsutsui et al. 2009), will be a useful approach for precise determination of evolutionary relationships (Duarte et al. 2010).

The availability of a suitable and precise reference genome is crucial in evolutionary studies to determine relationships among the species with higher accuracy. The haploid genome size of *M. indica* is estimated to be 439 Mbp by flow cytometric analysis (Arumuganathan and Earle 1991). The first draft genome for *M. indica* was assembled for the Indian cultivar Amrapali (Singh et al. 2014; Singh et al. 2018). The genomes of *M. indica* cv. Tommy Atkins (Bally et al. 2021), Kensington Pride (Dillon et al. 2016), Hong Xiang Ya (Li et al. 2020) and Alphonso (Wang et al. 2020) also have been sequenced using advanced sequencing platforms. A high-quality chromosome-level genome is available for the cultivar Alphonso (Wang et al. 2020). The genetics and genomics of chloroplasts have progressed rapidly with the advent of high-throughput sequencing technologies. Chloroplast genomes in higher plants are typically double-stranded and organized into conserved quadripartite structure, consisting of a pair of inverted repeats (IR) separated by small single copy region (SSC) and a large single copy region (LSC). The chloroplast genome size, although far smaller than most of the plant nuclear genomes, ranges from 120 kb to 160 kb (Odintsova and Yurina 2006) with 110 to 130 genes. Conflicts between the chloroplast and nuclear phylogenetic analysis provide valuable insights into speciation, hybridization and incomplete lineage sorting (Degnan and Rosenberg 2009; Joly et al. 2009). The first chloroplast genome sequence in genus *Mangifera* was reported for *M. indica (*Azim et al. 2014) and so far, assembled chloroplast genomes of only six out of 69 species (Jo et al. 2017; Niu et al. 2021) are available. In this study, we compared sequences of chloroplast genomes, and a selected set of common single-copy genes present in nuclear genome of 11 *Mangifera* species to analyse evolutionary relationships in the genus.

## 2. Materials and methods

### 2.1. Plant material and DNA extraction

A total of 13 samples belonging to 11 *Mangifera* species were selected for sequencing (Table 1). Leaf tissue of all *M. indica* varieties and *Mangifera* species, except *M. pajang* and *M. caesia*, are sourced from trees grafted onto *M. indica* cv. Kensington Pride rootstock at the Walkamin Research Station, Mareeba, (17°08̍ 02″S and 145°25 37″E), North Queensland, Australia. *M. pajang* and *M. caesia* were sourced from trees at Treefarm, El Arish (-17° 47’59.99”S and 146°00’0.00”E) and Fruit Forest Farm (www.fruitforestfarm.com.au, East Feluga, (17°53’46.0”S and 145°59’38.0”E), Queensland, Australia, respectively. Leaves after harvest were snap-frozen in liquid nitrogen, transported under dry ice and stored at -70℃ until processed for DNA extraction. Frozen leaf tissue was first coarsely ground using a mortar and pestle and then finely ground using the Qiagen Tissue lyser (Qiagen, USA). DNA extraction was carried out from fine pulverized mango leaf tissue samples using a cetyltrimethylammonium bromide method (Furtado 2014). The quality and quantity of DNA were assessed for acceptable absorbance ratios (ideal 1.8-2.0 at A260/280 and over 2.0 at 260/230) using Nanodrop Spectrophotometer (Thermo Fisher Scientific). DNA degradation and quantity were assessed by resolving sample and standard DNA by agarose gel electrophoresis (Thermo Fisher Scientific). The isolated DNA was subjected to next-generation short read sequencing (NGS) on an Illumina HiSeq 2000 platform at the Ramaciotti Centre for Genomics, University of New South Wales (UNSW), Australia to obtain sequence data with coverage of the genome that was not less than 20X (Table S1).

**Table 1:** Details of the 11 *Mangifera* species used in this study including country of origin, native distribution and important characteristics.

| Sub Genera | Section | Species/Taxon | Embryony | Country of Origin | Geographical distribution of the species | Ploidy Level (2n) | Prominent fruit/horticultural characteristics |
| --- | --- | --- | --- | --- | --- | --- | --- |
| *Mangifera* | *Mangifera* Ding Hou | *M. laurina* | Polyembryonic | Indonesia | Wild distribution: Myanmar, Vietnam, Malesia, Thailand. Cultivated in Borneo, Sumatra, Java, (Indonesia), and Philippines | 40 | Juicy, very acid and fibrous fruit: edible<br>Resistance to anthracnose (fruit skin)<br>Used as a rootstock for *M. indica* in Malaysia |
| *Limus* | *Deciduae* | *M. pajang* | Monoembryonic | Indonesia | Endemic to Borneo (Brunei Indonesia, Malaysia). Common in cultivation | Unknown | Fruit flesh: deep-orange-yellow, fibrous, acid to acid sweet, mildly fragrant: edible<br>Largest fruit in the genus |
| *Mangifera* | *Euantherae* Pierre | *M. caloneura* (Xoài Quéo) | Polyembryonic | Vietnam | Dry deciduous dipterocarp forests in Myanmar, Thailand, Laos, and Vietnam | 40 | Fruit with good aroma, sour taste: edible<br>Drought tolerance/ resistance to anthracnose (fruit skin) and gummosis |
| *Limus* | *Perrennis* | *M. odorata* | Polyembryonic | Malaysia | Never been observed in the wild<br>Origin is unknown, primarily cultivated in Guam, Philippines, Thailand, and Vietnam<br>Introduced to Indonesia, Malaysia, and Singapore | 40 | Firm, fibrous, sweet to acid-sweet, juicy fruit with a strong smell and turpentine flavour: edible<br>Resistance to anthracnose |
| *Mangifera* | *Mangifera* Ding Hou | *M. altissima* | Polyembryonic | Philippines | Native to Philippines, Indonesia, Papua New Guinea, Solomon Islands | Unknown | Smooth fibrous or non-fibrous fruit: edible |
| *Limus* | *Deciduae* | *M. caesia* | Monoembryonic | Indonesia | Natural distribution: Peninsular Malaysia, Borneo (Brunei Darussalam, Indonesia, Malaysia) and Sumatra (Indonesia)<br>Cultivated from Peninsular Thailand, Bali, Java and to the Philippines | 40 | Sweet flesh with strong fragrance in the fruit: edible |
| *Mangifera* | *Mangifera* Ding Hou | *M. zeylanica* | Monoembryonic | Sri Lanka | Endemic to Sri Lanka, not found under cultivation | Unknown | Very juicy fruit, fibrous, pleasant, sweet taste pulp: edible |
| *Mangifera* | *Mangifera* Ding Hou | *M. lalijiwa* | Polyembryonic | Indonesia | Indonesia (Java, Madura, Bali and probably in Sumatra). Very rare both in wild and in cultivation | Unknown | Small glossy yellowish fruit with acid-sweet taste:edible |
| *Mangifera* | *Mangifera* Ding Hou | *M. applanata* | Polyembryonic | Malesia | Native to Indonesia (Kalimantan and Sumatra) and Malaysia (Pahang, Sabah, and Sarawak)<br>Cultivated in some areas of Borneo | Unknown | Juicy, very acid fruit with turpentine and lemon taste: edible |
| *Mangifera* | *Mangifera* Ding Hou | *M. casturi* | Polyembryonic | Indonesia | Endemic to South Kalimantan, Indonesia.<br>Not known in the wild, only found in cultivation | Unknown | Small fruits, with an attractive purple color and a distinctive aroma: edible |
| *Mangifera* | *Mangifera* Ding Hou | *M. indica* cv. Kensington Pride | Polyembryonic | Australia | First planted in Bowen, North Queensland, Australia. Pre-Australian origin is unknown.<br>Grown throughout Australia | 40 | Soft and juicy flesh with moderate to little fibre, sweet with a characteristic flavour<br>excellent eating quality |
| *Mangifera* | *Mangifera* Ding Hou | *M. indica* cv. Alphonso | Monoembryonic | India | Prominently grown in India | 40 | Firm to soft flesh, low in fibre content, sweet with characteristic aroma, very pleasant taste: edible |
| *Mangifera* | *Mangifera* Ding Hou | *M. indica* cv. Tommy Atkins | Monoembryonic | USA (Florida) | Originated and most widely grown commercial cultivar in USA. | 40 | Fruit with firm, medium juicy, medium amount of fibre of good eating quality<br>Highly resistant to anthracnose disease |

### 2.2 Chloroplast genome Assembly

In addition to generating sequence data for the 11 *Mangifera* species, publicly available Illumina sequencing paired-end reads were downloaded from the National Centre for Biotechnology Information (NCBI) (http://www.ncbi.nlm.nih.gov/) for five *Mangifera* species namely *M. sylvatica*, *M. odorata*, *M. percisiformis*, *M. hiemalis* and *M. indica* cv. Tommy Atkins (Table S2). Chloroplast genomes for each species were assembled using two methods: a chloroplast assembly pipeline (CAP) described by (Moner et al. 2018) in CLC Genomic Workbench (CLC-GWB) software (CLC Genomics Workbench 20.0, http://www.clcbio.com) and “Get Organelle” pipeline (http://github.com/Kinggerm/GetOrganelle) (Jin et al. 2020). Raw reads for all the species were imported to CLC-GWB and trimmed using the quality score limits of 0.01 (Phred score equivalent to >20) at the sequence length of 1000. The CAP processed two approaches to assemble the chloroplast genome, a reference-guided mapping approach and a de-novo assembly approach. For the reference guided mapping, *M. indica* cv. Tommy Atkins chloroplast genome (Accession: NC_035239.1) (Rabah et al. 2017), available from NCBI, was imported to CLC-GWB and used as the reference to assemble the chloroplast genomes. The two chloroplast sequences generated using the two approaches of the CAP for each species were aligned in Geneious 2022.2.2 software (www.geneious.com) and Clone Manager Professional 9 to identify mismatches. Manual curation of mismatches involved observing the reads mappings at the position of the mismatch. De-novo assembled chloroplast genomes from Get Organelle pipeline were checked in Bandage v. 0.8.1(Wick et al. 2015) to visualize the completeness of the assembled genomes. The final chloroplast genome assembled from CAP and Get Organelle pipeline were compared for mismatches and further manual curation, ensuring high-quality chloroplast genomes were assembled for all the species.

### 2.3. Chloroplast genome annotation and identification of single nucleotide polymorphisms (SNPs), insertions and deletions (INDELs)

Genome annotations for assembled chloroplast genomes were performed using GeSeq online tool (https://chlorobox.mpimp-golm.mpg.de/geseq.html) and *M. indica* cv. Tommy Atkins (Accession: NC_035239.1) was used as the reference genome. Based on the evolutionary relationships observed in chloroplast genome-based phylogenetic analysis, closely related species within the main clades and subclades were compared to determine their evolutionary relationships. Chloroplast genomes of the species were subjected to pairwise alignment in Geneious using the MAFFT alignment tool and the number INDELs, substitutions and SNPs present between the sequences were identified.

### 2.4. Nuclear gene sequence assembly

Here, we used a list of single copy nuclear genes. Details of the genes were not available for *Mangifera* species. Therefore, *Citrus sinensis*, the closest relative of *M. indica* for which the details of single copy nuclear genes were available (Li et al. 2017) was used as the reference, to extract corresponding single-copy genes in mango. Single copy genes (107) in *C. sinensis* were mapped against the coding DNA sequences/gene models of *M. indica* cv. Alphonso (Wang et al. 2020) in CLC-GWB. Out of 107, 47 were identified as single-copy genes in *M. indica*. Then, trimmed paired-end illumina reads of each species were mapped against 47 single-copy genes of *M. indica,* and consensus gene sequences were extracted. All 47 genes were identified as single-copy genes in *Mangifera* species. Also, the same 47 gene sequences were extracted from *A. occidentale.* Details of the single copy genes in mango are indicated in Table S3.

### 2.5. Phylogenetic analysis

Phylogenetic analysis of *Mangifera* species was undertaken using the chloroplast genome sequences and also for the single-copy nuclear gene sequences. Corresponding sequences of *A. occidentale* were used as an outgroup. Apart from the 13 samples sequenced in this study, five species for which chloroplast genomes and nuclear gene sequences generated by downloading sequence data from NCBI were also included.

#### 2.5.1. Chloroplast genome-based phylogenetic analysis

For chloroplast genomes-based phylogenetic analysis, Sequences were imported to the Geneious 2022.2.2 software (www.geneious.com) and aligned by multiple sequence alignment using MAFFT (MAFFT v7.490) alignment tool (Katoh and Standley 2013). Two methods were used for phylogenetic analysis: Maximum likelihood (ML) method and Bayesian inference (BI) method. jMfodelTest v2.1.4 (Darriba et al. 2012) was used to select the best fitting nucleotide substitution model using Cyberinfrastructure for Phylogenetic Research (CIPRES) Science Gateway (http://www.phylo.org/) (Table S4). ML analysis was performed in RaXML GUI 2.0 (v 2.0.10) (Stamatakis 2014) with 1000 bootstrap replicates under Akaike information criterion. Bayesian analysis was carried out in Geneious software using MrBayes v. 3.2 (Ronquist et al. 2012) under the Bayesian information criterion (Table S4). iTOL v.6 tool (https://itol.embl.de/) (Letunic and Bork, 2021) was used to visualize and edit the phylogenies. Using posterior probability (PP) and bootstrap support (BS) to evaluate the supports of the phylogenetic tree implemented under BI and ML methods respectively, final trees obtained from both approaches were compared and tree topologies were validated.

#### 2.5.2. Nuclear gene-based phylogenetic analysis

For nuclear gene sequences, phylogenetic trees were generated using two approaches: gene concatenation and coalescent approach to analyse any topological incongruence and for a better understanding of evolutionary relationships among species.

##### Concatenation approach

All 47 genes of were concatenated in the same order to get a one long sequence per species. Sequences for all the species were imported to the Geneious 2022.2.2 software (www.geneious.com) and aligned by MAFFT alignment. Phylogenetic trees were constructed using ML method and BI methods after selecting the best fitting nucleotide substitution model by running jMfodelTest v2.1.4 (Darriba et al. 2012) (Table S4). ML analysis was performed in RAxML (version 8) (Stamatakis 2014) with 1000 bootstrap replicates, and Bayesian analysis was carried out in Geneious software using MrBayes v. 3.2 (Ronquist et al. 2012) (Table S4). iTOL v.6 tool (https://itol.embl.de/) (Letunic and Bork 2021) was used to visualize and edit the phylogenetic trees. By using PP and BS values to evaluate the supports of the phylogenetic tree implemented under BI and ML methods respectively, final trees obtained from both approaches were compared and tree topologies were validated.

##### Coalescent approach

For coalescent approach, single ML gene trees were constructed by ML method using RAxML (version 8) (Stamatakis 2014). We searched for the best-scoring ML tree using a GTR+GAMMA model with 1000 bootstrap replicates. Low support branches (BS<10%) in gene trees were collapsed to minimize potential impacts of gene tree error for species tree reconstruction. Then, these gene trees were used to construct a coalescent-based species tree using ASTRAL-III (Zhang et al. 2018)

## 3. Results

### 3.1. Chloroplast genome assembly and annotation

Illumina sequencing conducted for the 13 samples belonging to 11 *Mangifera* species in this study resulted in 60,699,616 to 181,601,786 raw reads with 150bp mean read length. Trimmed paired-end reads at 0.01 quality limits (Phred score > 20) ranged between 59,763,897 and 171,303,402 reads. The data size of the trimmed reads of all 13 samples corresponded to over 20x of the genome size, therefore all were selected for the chloroplast assembly (Table S1). Raw reads downloaded from NCBI for five *Mangifera* species (*M. odorata*, *M. sylvatica*, *M. percisiformis*, *M. hiemalis* and *M. indica* cv. Tommy Atkins) had a total of 99,649,506 to 127,708,722 reads which ranged from 94,450,606 to 117,445,811 after trimming at 0.01 quality limits. For all the species, the mean coverage was higher than 20x genome size, which enabled them to be included in the analysis (Table S2). The chloroplast genome and respective raw reads were also available for the species *M. longipes* (synonym*: M. laurina*), but the average coverage after quality trimming was less than 20x, and therefore this was not included for the analysis. The Get Organelle pipeline resulted in two output files for the chloroplast genome for each species/ genotype, due to possibility of the SSC occurring in both orientations in the chloroplast genomes in plants. Therefore, the two chloroplast sequences for each species were aligned with the reference (*M. indica*; Accession: NC_035239.1) in Clone Manager Professional 9 to select the sequence with the widely accepted SSC orientation (5’LSC3’:5’IR13’:5SSC3’:3’IR25’). The size of the chloroplast genomes of 13 wild *Mangifera* species and three cultivars of *M. indica* ranged from 151,752bp to158,965bp of which the smallest and the largest genomes were recorded for *M. caesia* and *M. laurina* Lombok, respectively (Table 2). The typical quadripartite structure of the chloroplast genome was recorded in all 14 *Mangifera* species and the lengths of LSC, SSC, and IR regions ranged between 86,507 to 98,334 bp, 18,319 to 19,064 bp and 17,177 to 26,412 bp, respectively where overall guanine – cytosine content (GC content) ranged from 37.6 to 37.9 %. The chloroplast genomes for all species had the same number of total genes (115), rRNA (4) and tRNA (31) and protein encoding genes (80) (Table 2). Although the size of the chloroplast genomes varies across the *Mangifera* species, three cultivars of *M. indica* had identical chloroplast genomes. Two *M. odorata* accessions had slightly different chloroplast genome sizes, where the accession we sequenced had a genome size of 158,889bp while for the sample for which the data was downloaded from NCBI (*M. odorata*\*) had a genome size of 158,883 bp, representing a 5 bp difference. The length difference was due to two deletions revealed in *M. odorata*\*, one located in a non- coding region of LSC while the other located in the intron1 region of the *PetD* gene of LSC. The chloroplast sequence of *M. indica* cultivar Kensington Pride (Fig. 1) is a representation of the chloroplast sequence of the 14 *Mangifera* species which have the same number of genes although there are differences in total chloroplast size and the sizes of the LSC, IR1 and IR2 and the SSC regions.

**Fig. 1.**
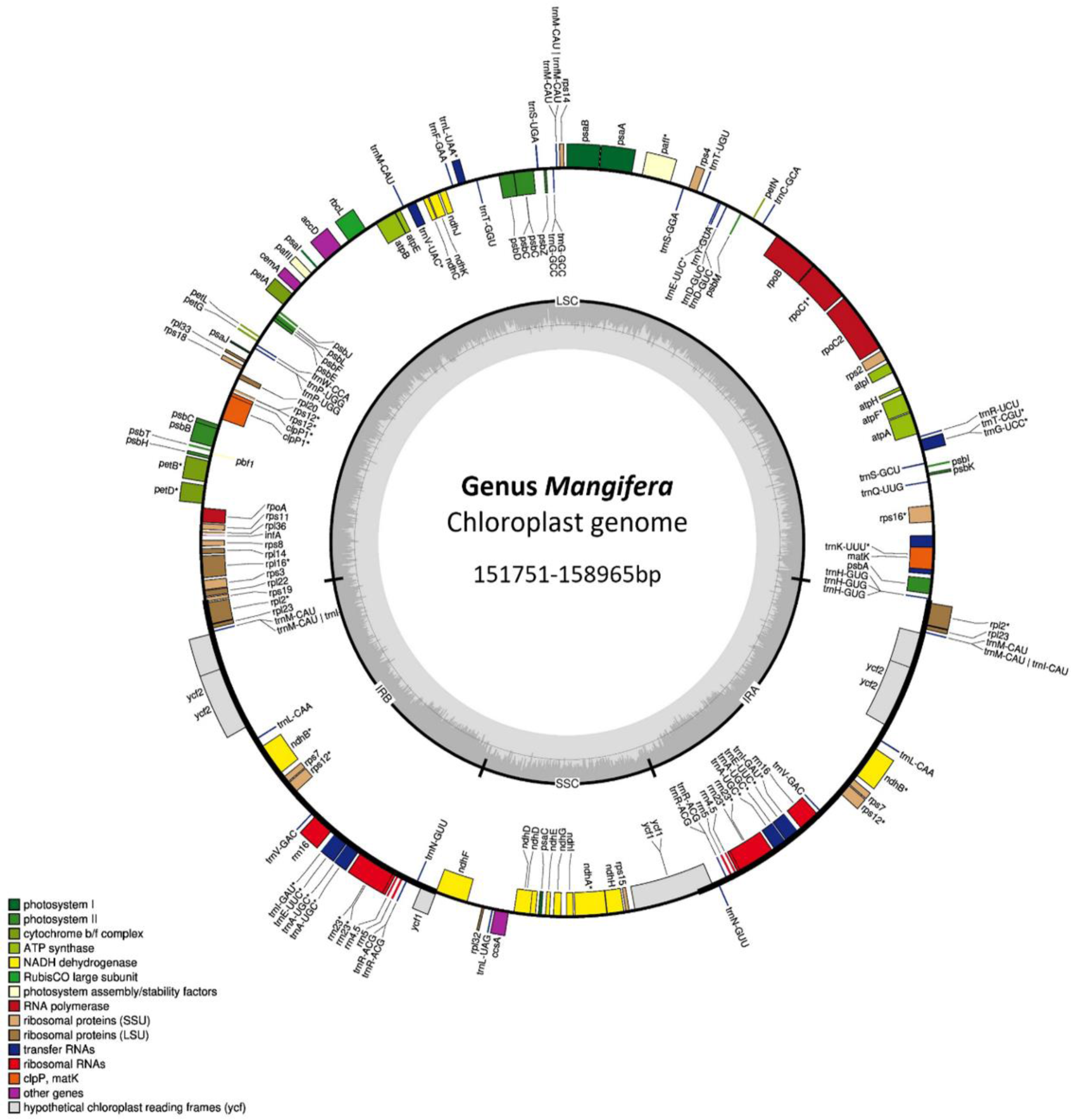
**Genome map of chloroplasts in the genus *Mangifera.*** The genome size of the 14 *Mangifera* species ranges from 151,752 to 158,965 bp for *M. caesia* and *M. laurina* Lombok, respectively. In the outer most circle, the black thick border/line indicates Inverted Repeat Regions (IR) whereas the thin lines indicate Large Single Copy (LSC) and the Small Single Copy (SSC). Genes inside the circle are transcribed in the clockwise direction whereas the genes outside the circle are transcribed in the counter-clockwise direction. Different colours are given for the genes with respect to their functions. The darker grey in the inner circle corresponds to GC content, whereas the lighter grey corresponds to AT content

**Table 2:** Details on genome annotation results of *Mangifera* species.

| Species/Genotype | Genome Size (bp) | Overall GC content (%) | LSC (bp) | SSC (bp) | IR (bp) | Total number of genes | Total number of protein coding genes | Total number of tRNAs | Total number of rRNAs |
| --- | --- | --- | --- | --- | --- | --- | --- | --- | --- |
| *M. laurina* | 158,965 | 37.8 | 87,714 | 18,427 | 26,412 | 115 | 80 | 31 | 4 |
| *M. pajang* | 158,830 | 37.8 | 87,654 | 18,424 | 26,376 | 115 | 80 | 31 | 4 |
| *M. caloneura* (Xoài Quéo) | 157,780 | 37.9 | 86,672 | 18,350 | 26,379 | 115 | 80 | 31 | 4 |
| *M. odorata* | 158,889 | 37.8 | 87,708 | 18,427 | 26,377 | 115 | 80 | 31 | 4 |
| *M. altissima* | 157,780 | 37.9 | 86,673 | 18,349 | 26,379 | 115 | 80 | 31 | 4 |
| *M. caesia* | 151,752 | 37.6 | 98,334 | 19,064 | 17,177 | 115 | 80 | 31 | 4 |
| *M. zeylanica* | 157,604 | 37.9 | 86,507 | 18,319 | 26,389 | 115 | 80 | 31 | 4 |
| *M. lalijiwa* | 157,779 | 37.9 | 86,672 | 18,349 | 26,379 | 115 | 80 | 31 | 4 |
| *M. applanata* | 157,779 | 37.9 | 86,672 | 18,349 | 26,379 | 115 | 80 | 31 | 4 |
| *M. casturi* | 158,942 | 37.8 | 87,733 | 18,425 | 26,392 | 115 | 80 | 31 | 4 |
| *M. sylvatica* * | 157,781 | 37.9 | 86,712 | 18,347 | 26,361 | 115 | 80 | 31 | 4 |
| *M. odorata* * | 158,883 | 37.8 | 87,702 | 18,427 | 26,377 | 115 | 80 | 31 | 4 |
| *M. persiciformis* * | 158,952 | 37.8 | 87,566 | 18,536 | 26,368 | 115 | 80 | 31 | 4 |
| *M. hiemalis* * | 158,838 | 37.8 | 87,681 | 18,535 | 26,368 | 115 | 80 | 31 | 4 |
| *M. indica* cv. Kensington Pride | 157,780 | 37.9 | 86,673 | 18,349 | 26,379 | 115 | 80 | 31 | 4 |
| *M. indica* cv. Alphonso | 157,780 | 37.9 | 86,673 | 18,349 | 26,379 | 115 | 80 | 31 | 4 |
| *M. indica* cv. Tommy Atkins* | 157,780 | 37.9 | 86,673 | 18,349 | 26,379 | 115 | 80 | 31 | 4 |

- Note. GC, guanine or cytosine; IR, inverted repeats; LSC, large single copy; SSC, small single copy - \* *Mangifera* species for which chloroplast genomes were assembled using raw data downloaded from NCBI

### 3.2. Chloroplast phylogeny and identification of SNPs and INDELs

A multiple chloroplast sequence alignment conducted using *A. occidentale* as the outgroup followed by phylogenetic tree construction resulted in a ML tree and a Bayesian tree with same tree topology. BS and PP values of the final tree are presented in Fig. 2. The model of nucleotide substitutions for ML analysis was GTR+G whereas for the Bayesian analysis, it was TPM1uf+G. The tree developed with the ML approach showed a BS of 100 at most of the nodes and PP of one in all the nodes. In the whole plastome tree, three main clades were identified. First, 14 *Mangifera* species were clustered into two distinct clades in which only *M. caesia* belonging to section *Dissidue* in the subgenus *Limus* was placed in first clade (Clade A). Other 13 species were grouped into a separate clade indicating their evolutionary distinct relationship to *M. caesia,* which were then clustered into two sub clades (clade B and clade C). Clade B included a total of six species which belong to different categories in the classification. *M. pajang* and *M. odorata* belong to sub genus *Limus* while *M. casturi* and *M. laurina* belong to subgenus *Mangifera*. *M. percisiformis* and *M. hiemalis* are two species placed under uncertain position in the classification. Within clade B, species in subgenera *Mangifera* (Clade BI), *Limus* (Clade BII), and species being classified in an uncertain position (Clade BIII) have localized into well supported distinct clades (BS=100, PP=1). The species belong to subgenera *Mangifera* and *Limus* and were sister to each other and both together have become a sister clade to species placed in uncertain position in the classification. Clade C had species belonging only to the sub genus *Mangifera* which was further divided into three subclades (clades CI, CII and CIII). Interestingly, four wild species (*M. lalijiwa*, *M. applanata, M. altissima* and *M. caloneura*) were clustered with three cultivars of domesticated mango (*M. indica*) (Clade CI). Although species belong to section *Mangifera* and *Euantherae* are characterised by the presence of one and multiple fertile stamens respectively, *M. caloneura* in section *Euantherae* was clustered with species belonging to section *Mangifera*. Furthermore, *M. sylvatica* and *M. zeylanica*; two species within the clade C were separately clustered into distinct clades, CII and CIII respectively.

**Fig. 2.**
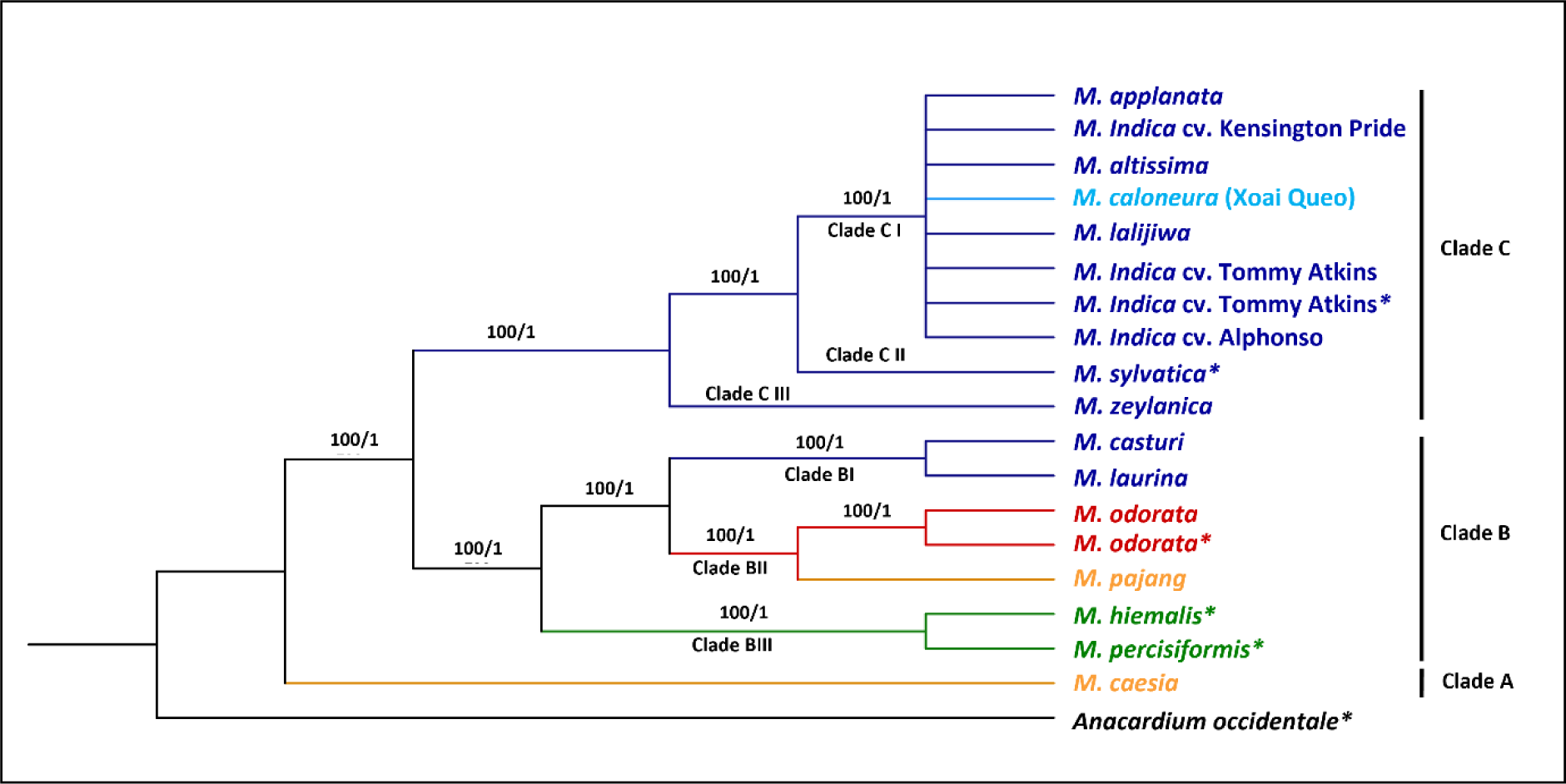
The phylogenetic tree developed for *Mangifera* species based on whole chloroplast genomes. Phylogenetic tree of 17 accessions belong to 14 species with *A. occidentale* used as the outgroup. Trees were generated using Maximum Likelihood (ML) and Bayesian inference (BI) method. Numbers associated with the branches are ML bootstrap value (/100) and BI posterior probabilities (/1). Dark Blue: Sub genus *Mangifera*, Section *Mangifera*, Light blue: Sub genus *Mangifera*, Section *Euantherae,* Red: Sub genus: *Limus*, Section *Perrennis,* Yellow: sub genus: *Limus*, Section: *Deciduae.* * *Mangifera* species for which chloroplast genomes were assembled using raw data downloaded from NCBI

*M. sylvatica* is the sister to domesticated clade (clade CI) and *M. zeylanica* (Clade CIII) has become sister to the clade includes CI and CII. Therefore, the phylogeny based on whole chloroplast genome clustered species belong to different groups inferring the close genetic and evolutionary relationships of their chloroplast genomes (Fig. 2).

Assembled chloroplast genomes were imported to Geneious software to conduct pairwise alignment to identify number and types of variants present between the species clustered as sister taxa in the two main clades (clade B and C) of the chloroplast phylogeny. A total of 116 variants were observed between *M. laurina* and *M. casturi* while 75 variants were found between *M. odorata* and *M. pajang* (Table 3). The two species, *M. percisiformis* and *M. hiemalis,* for which we assembled the chloroplast genomes from raw read data available in NCBI, differed by 45 variants revealing the close evolutionary relationships among these sister taxa in clade BIII. In clade CI, pairwise comparison of wild species with *M. indica* cv Kensington Pride showed that, *M. altissima* and *M. indica* had identical chloroplast sequences. Moreover, *M. lalijiwa* and *M. applanata* also had an identical chloroplast genome which only differed from *M. indica* by having a single nucleotide deletion located in a non-coding region in LSC. Furthermore, despite reporting distinct morphological characteristics from *M. indica*, *M. caloneura* only had one single nucleotide insertion and one single nucleotide deletion in non-coding regions compared to *M. indica*. Diversity within the chloroplast genomes clustered in clade CI was very low.

**Table 3:**
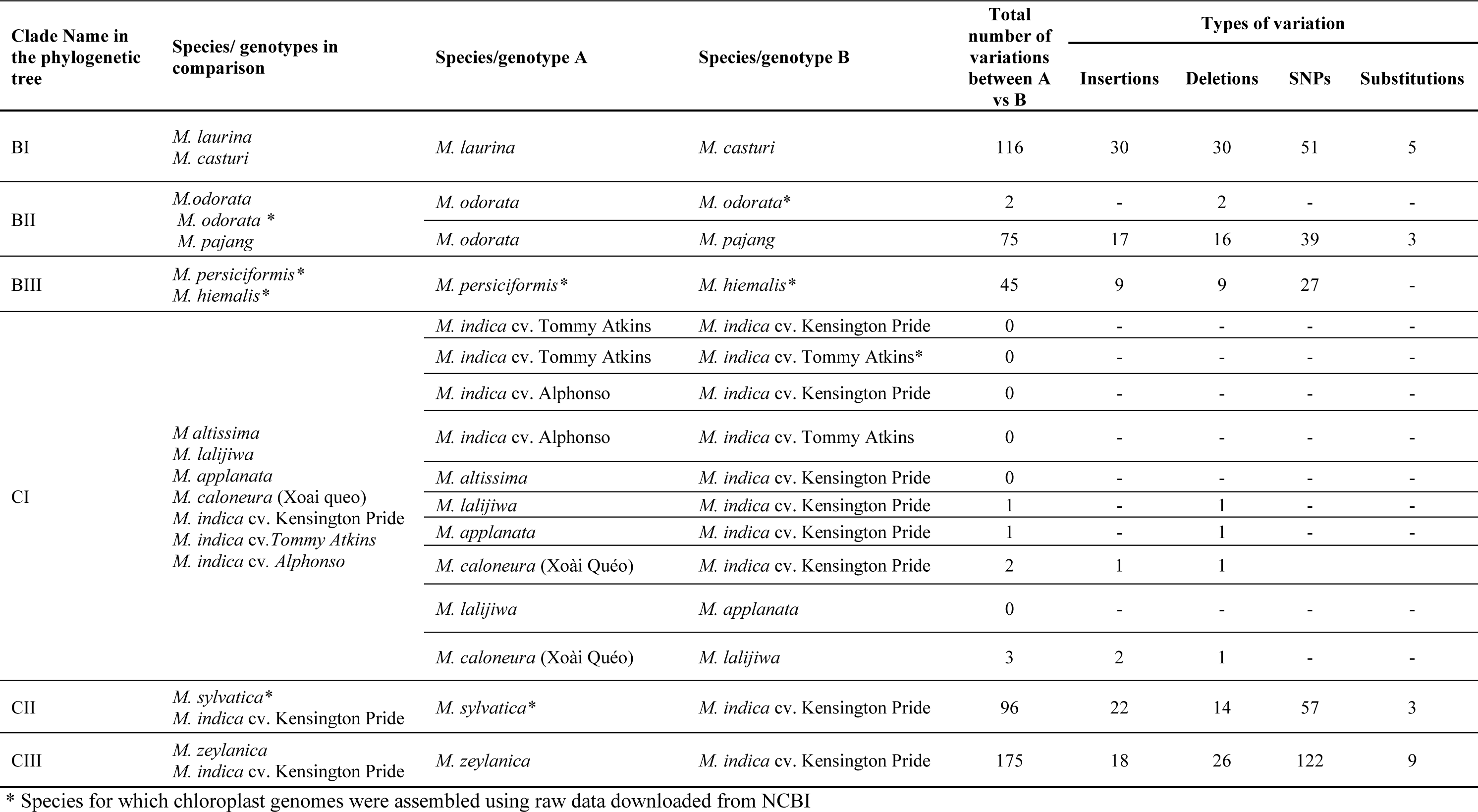
Details of INDELs, SNPs and substitutions identified with respect to clustering pattern in chloroplast phylogeny.

### 3.3. Nuclear gene phylogeny

#### 3.3.1. Concatenation-based nuclear phylogeny

The same approach was used to construct a nuclear phylogeny with concatenation-based approach as was applied in constructing the chloroplast phylogeny. *A. occidentale* was used as the outgroup. A total of 47 common single copy nuclear genes out of 107 (Li et al. 2017) were identified and selected for *Mangifera* species. The multiple sequence alignment was 71,881 bp in length and ML and Bayesian trees resulted in the same tree topology. The final tree with BS values and PP values is presented in Fig. 3a. Although some of the nodes showed less BS support values, all the nodes were supported with high PP values. The model of nucleotide substitutions for ML analysis was GTR+I+G whereas TPM1+I+G was used for the Bayesian analysis.

**Fig. 3.**
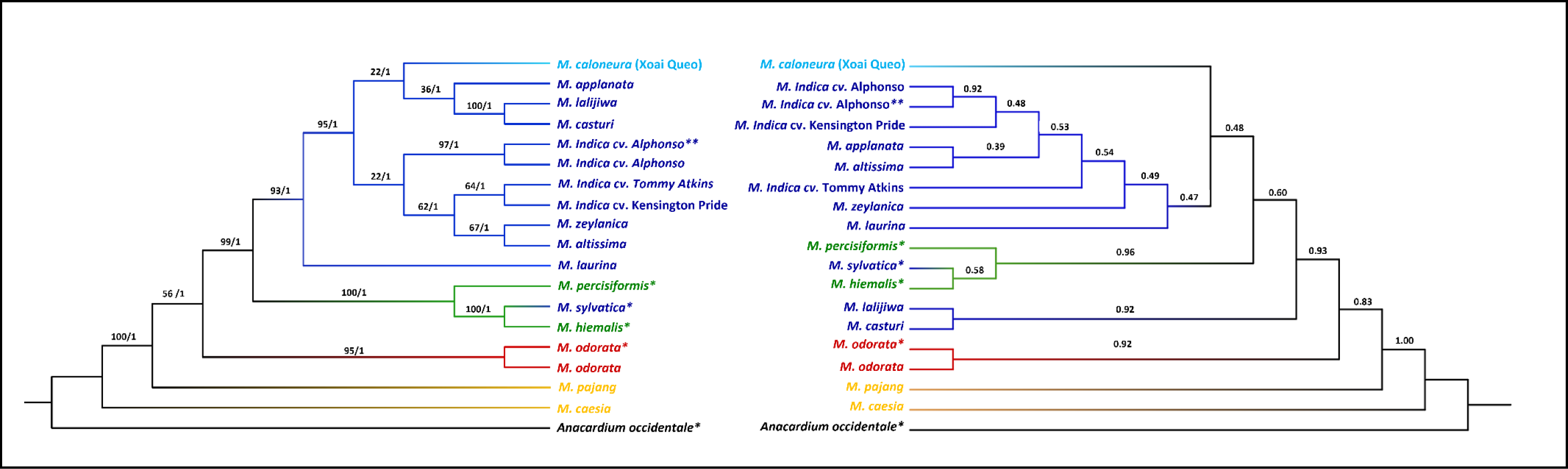
The phylogenetic tree developed for *Mangifera* species based on a selected set of nuclear genes using a) concatenation and b) coalescence-based methods. Phylogenetic tree of 17 accessions with *A. occidentale* used as the outgroup. Concatenation-based trees were generated using Maximum Likelihood (ML) and Bayesian inference (BI) methods and consensus tree is shown in the figure. Numbers associated with the branches are ML bootstrap value (/100) and BI posterior probabilities (/1). In the coalescence-based tree (ASTRAL tree), numbers associated with branches are local posterior probability values (/1). Dark Blue: Sub genus *Mangifera*, Section *Mangifera*, Light blue: Sub genus *Mangifera*, Section *Euantherae,* Red: Sub genus: *Limus*, Section *Perrennis,* Yellow: sub genus: *Limus*, Section: *Deciduae.* * Species for which nuclear genes were extracted using raw data downloaded from NCBI. ** *M. indica* cultivar from which gene models were downloaded from NCBI and used to create local database in CLC-GWB for the selection of single copy nuclear genes in *M. indica*

Except for *M. sylvatica,* the other eight species belonging to subgenus *Mangifera* were clustered into one main distinct clade. *M. lalijiwa*, *M. applanata* and *M. caloneura* with *M. casturi* were clustered into one clade and *M. altissima*, *M. zeylanica*, and *M. indica* cultivars were clustered into another clade within the main clade. *M. applanata* and *M. lalijiwa* were sister taxa to each other. Furthermore, two *M. indica* cultivars (Kensington Pride and Tommy Atkins) showed closer genetic relationship to *M. zeylanica* and *M. altissima* revealing close evolutionary relationship of the two wild species to domesticated mango. Moreover, although *M. sylvatica* was closely related to species in domesticated clade in chloroplast phylogeny, it was clustered with the two species placed in uncertain position in the classification in the nuclear phylogeny where it was more closely related to *M. hiemalis* than to *M. percisiformis*.

Furthermore, both chloroplast and concatenation-based nuclear phylogenies revealed that *M. caesia* is evolutionarily distant from the rest of the *Mangifera* species (Fig. 2 and 3a). Grouping of species in both chloroplast genome and nuclear genes-based analysis does not completely concur with the accepted classification described by (Kostermans and Bompard 1993) for genus *Mangifera.* Incongruence in tree topologies could be seen between the phylogenies developed based on the whole plastome genome and the nuclear genes.

#### 3.3.2. Coalescence-based nuclear phylogeny

Previously proposed hybrids were included in our dataset and possibility of hybridization events also were observed for some species when compared chloroplast and concatenation-based nuclear phylogenies. Therefore, to further analyse the phylogenetic relationships among *Mangifera* species with respect to nuclear genes, coalescence approach was utilised to develop individual nuclear gene trees thereby to develop a species tree. Individual gene trees were analysed to see close evolutionary relationships among species.

In coalescence-based species tree, local posterior probability support (LPP) values are indicated in the branches (/1). In both concatenation and coalescence based nuclear phylogenies, species belonging to subgenus *Mangifera,* and species placed in uncertain position in the classification were clustered in one clade with high support values (BS=99, PP=1, LPP=0.93) (Fig. 3). Within this clade, pattern of clustering into sub-clades was different for some species between the two nuclear phylogenies but BS and LPP support values also were low for some sub-clades. In both concatenation and coalescence-based phylogenies, *M.casturi* and *M.lalijiwa* are clustered together as sister taxa (BS=100, PP1, LPP= 0.92) and *M. hiemalis*, *M. sylvatica* and *M. percisiformis* were clustered in to one sub-clade (BS=100, PP=1, LPP=0.96). But in coalescence-based tree, *M. hiemalis*, *M. sylvatica* and *M. percisiformis* were closely related to six other species belong to subgenus *Mangifera* (*M. altissima*, *M. applanata*, *M.indica*, *M. caloneura*, *M.zeylanica* and *M. laurina*) than *M. casturi* and *M. lalijiwa* while these eight species in subgenus *Mangifera* were clustered together into one clade in concatenation-based tree.

*M.odorata* is a proposed hybrid between *M. indica* and *M. foetida* and *M. casturi* is a proposed hybrid between *M. indica* and *M. quadrifida*. Within our dataset, only one of the parents are available for these hybrids. Although it is not possible to validate the hybridity due to absence of one of the parents, we analysed individual gene trees to support the hybridity by recording the number of gene trees where the hybrids were clustered with the parent available in our dataset (*M. indica*). Out of 47 gene trees, *M. odorata* was clustered with *M. indica* as sister taxa in only four gene trees and they were not supported with high BS values (Table 4, Figure S1). Similarly, *M. casturi* was also clustered with *M. indica* as sister taxa in four gene trees only in which BS support were weak in three gene trees for this clade (Table 4, Figure S1). We observed close evolutionary relationship between *M. zeylanica* and *M. indica* and also between *M. hiemalis and M. sylvatica* in nuclear phylogenies. Therefore, we assumed that *M. zeylanica* might have undergone domestication and *M. sylvatica* may have cross hybridised with *M. hiemalis* during evolution of these species. Analysing individual gene trees revealed that *M. zeylanica* was clustered with *M. indica* as sister taxa in 14 gene trees and *M. sylvatica* was clustered with and *M. hiemalis* as sister taxa in 12 gene trees. Some individual gene trees for *M. zeylanica* and *M. sylvatica* showed less BS support when clustering with *M. indica* and *M. hiemalis* respectively (Table 4, Figure S1).

**Table 4:** Gene trees indicating clustering of suggested hybrids/ wild relatives with their proposed parents as sister taxa.

| Gene name | Gene Tree No | Species proposed to be hybrid |  | Species supposed to be undergone domestication/hybridization |  |
| --- | --- | --- | --- | --- | --- |
|  |  | *M.odorata* | *M. casturi* | *M.zeylanica* | *M.sylvatica* |
|  |  | Trees in which *M.odorata* clusters with *M.indica* in the same clade as sister taxa | Trees in which *M.casturi* clusters with *M.indica* in the same clade as sister taxa | Trees in which *M.zeylanica* clusters with *M.indica* in the same clade as sister taxa | Trees in which *M.sylvatica* clusters with *M.hiemalis* in the same clade as sister taxa |
| 4-alpha-glucanotransferase | 1 | × | × | √ | × |
| A49-like RNA polymerase I associated factor | 2 | × | × | √ | × |
| Acyl-CoA dehydrogenase, C-terminal domain | 3 | × | × | × | × |
| Alphabeta hydrolase family | 4 | × | × | √ | × |
| Aminomethyltransferase folate-binding domain | 5 | × | × | √ | × |
| Aminopeptidase I zinc metalloprotease (M18) | 6 | × | × | × | × |
| ArgJ family | 7 | × | × | × | × |
| Armadillobeta-catenin-like repeat | 8 | × | × | × | × |
| Brix domain | 9 | √ | × | × | × |
| Cactus-binding C-terminus of cactin protein | 10 | × | × | × | × |
| Carbon-nitrogen hydrolase | 11 | √ | × | × | × |
| CobWHypBUREG, nucleotide-binding domain | 12 | × | √ | × | × |
| Cohesin loading factor | 13 | × | × | × | √ |
| CreatinaseProlidase N-terminal domain | 14 | × | × | × | × |
| Cyclophilin type peptidyl-prolyl cis-trans isomeraseCLD | 15 | × | × | × | × |
| Cytochrome b5-like HemeSteroid binding domain | 16 | × | × | × | × |
| DDRGK domain | 17 | × | × | √ | × |
| Dienelactone hydrolase family | 18 | × | × | × | √ |
| Divergent CRALTRIO domain | 19 | × | × | √ | √ |
| Dual specificity phosphatase, catalytic domain | 20 | × | × | √ | √ |
| Dynamain family (2) | 21 | × | × | × | × |
| ER membrane protein complex subunit 1, C-terminal | 22 | × | × | × | × |
| Eukaryotic protein of unknown function (DUF866) | 23 | × | √ | × | √ |
| FAD binding domain | 24 | × | × | × | × |
| Glucose-6-phosphate isomerase | 25 | × | × | × | × |
| Glucosidase II beta subunit-like protein | 26 | × | × | × | √ |
| Glycosyl hydrolases family 31 | 27 | × | × | × | √ |
| GlyoxalaseBleomycin resistance proteinDioxygenase superfamily | 28 | × | × | × | × |
| Hydroxyacylglutathione hydrolase C-terminus | 29 | × | × | √ | × |
| Methyltransferase domain | 30 | × | × | × | × |
| NmrA-like family | 31 | × | × | √ | √ |
| N-terminal domain of lipoyl synthase of Radical_SAM family | 32 | × | × | × | × |
| PAP2 superfamily C-terminal | 33 | × | √ | × | × |
| Prolyl oligopeptidase, N-terminal beta-propeller domain | 34 | × | × | × | × |
| Putative tRNA binding domain | 35 | × | × | × | × |
| Pyruvate phosphate dikinase, AMPATP-binding domain | 36 | × | × | √ | √ |
| Redoxin | 37 | √ | × | × | × |
| Ribosomal protein L13 | 38 | × | × | √ | √ |
| RNA recognition motif. (a.k.a. RRM, RBD, or RNP domain) | 39 | × | × | × | √ |
| Rubryerythrin | 40 | × | × | × | × |
| snoRNA binding domain, fibrillarin | 41 | × | × | √ | √ |
| Sodium Bile acid symporter family | 42 | × | × | √ | × |
| TFIIS helical bundle-like domain | 43 | × | × | √ | × |
| Transcription factor TFIIB repeat | 44 | √ | × | × | × |
| tRNA synthetase class II core domain (G, H, P, S and T) | 45 | × | × | × | × |
| Uncharacterized ACR, YdiUUPF0061 family | 46 | × | √ | × | × |
| WLM domain | 47 | × | × | × | × |

## 4. Discussion

Determination of phylogenetic relationships among crop species provides basic information for predicting their evolutionary history, taxonomical classification, evaluating their diversity and importance in plant breeding (Zhang et al. 2012). Although genetic analysis of plants has improved rapidly with advanced sequencing technology, many phylogenetic studies in the genus *Mangifera* have relied on a set of molecular markers such as amplified fragment length polymorphisms (AFLP), rapid amplified polymorphic DNA (RAPD) and simple sequence repeats (SSR) and the sequencing of limited numbers of targeted regions in the chloroplast genome (*maturase K*, *trnL-F* spacer regions) and nuclear ribosomal DNA (internal transcribed spacer/*ITS* region) (Eiadthong et al. 1999; Fitmawati et al. 2017; Fitmawati 2016; Hartana 2010; Hidayat et al. 2011; Schnell and Knight Jr 1992; Yonemori et al. 2002). This is the first comparative analysis of phylogenetic relationship within the genus using both whole chloroplast genomes and multiple single-copy nuclear genes.

Chloroplast genomes for seven species were assembled for the first time in this study for *M. pajang*, *M. altissima*, *M. caesia*, *M. lalijiwa*, *M.* z*eylanica*, *M. appalanta* and *M. casturi*. Different pipelines and programs such as SPAdes (Bankevich et al. 2012), SOAPdenovo2 (Luo et al. 2012), ORG.Asm (Coissac et al. 2016), IOGA (Bakker et al. 2016), Fast-Plast (Afinit 2017), Organelle_PBA (Soorni et al. 2017), NOVOPlasty (Dierckxsens et al. 2017), chloroExtractor (Ankenbrand et al. 2018), CAP (Moner et al. 2018) and Get Organelle toolkit (Jin et al. 2020) are available to assemble organelle genomes. Here, we have used CAP and Get Organelle pipeline to assemble plastome genomes independently for each species. The two approaches used in CAP (reference-guided mapping and de-novo assembly) eliminate many errors in genomes developed from each approach giving a highly accurate final chloroplast genome. The Get Organelle pipeline; a fast and versatile toolkit used to assemble organelle genomes via de novo approach is also capable of generating all possible arrangements of the chloroplast genome present because of flip-flop configurations or other isomers mediated by repeats (Jin et al. 2020). Therefore, a comparison of chloroplast genomes generated from CAP and the Get Organelle pipeline validated the development of highly accurate final chloroplast genomes for all the species. Annotation of chloroplast genome with Ge Seq software discovered a total of 115 genes including 80 protein-coding genes, 31 tRNA genes, and four rRNA genes for all the species used in this study. More genes have been annotated in this analysis compared to previous studies, in which (Zhang et al. 2020) reported a total of 112 genes (78 protein-coding genes, 30tRNA genes, 4 rRNA genes) for *M. sylvatica* while 113 genes (79 protein-coding genes, 30tRNA genes, 4 rRNA genes) for *M. sylvatica*, *M. odorata*, *M. longipes*, *M. percisiformis, M. hiemalis* and *M. indica* were reported by (Niu et al. 2021).

Phylogenetic relationships within genus *Mangifera* showed topological incongruence for some species with respect to whole chloroplast and nuclear genes trees which may be caused by introgressive hybridization, allopolyploidy or incomplete lineage sorting. Reproductive compatibility between different species allows native cytoplasm of a species to be easily replaced by another through hybridization which has been detected both in animals (mitochondrial capture) (Liu et al. 2016; Rebbeck et al. 2011) and plants (chloroplast capture) (Rieseberg 1995; Rieseberg and Soltis 1991). In plants, chloroplast capture events have been reported in many plant families including *Maleae, Rosaceae* (Liu et al. 2020), *Nothofagaceae* (Acosta and Premoli 2010) *Scrophulariaceae* (Wolfe and Elisens 1995), *<u>Apiaceae</u>* (Yi et al. 2015), *Poaceae* (Ananda et al. 2021; Moner et al. 2020), *Rubiaceae* (Charr et al. 2020; Guyeux et al. 2019) and *<u>Myrtaceae</u>* (Healey et al. 2018). Hybridization followed by recurrent backcrossing have explained discrepancies between chloroplast and nuclear gene-based phylogenies in diverse families of plants (Liu et al. 2017; Smith and Sytsma 1990; Stegemann et al. 2012; Tsitrone et al. 2003). In mango, evidence for inter-specific reproductive compatibility was reported for *M. indica* and *M. laurina*. A cross between *M. indica* and *M. laurina* have produced 60 successful hybrids (Bally et al. 2010). Hybrid origins were reported for *M. odorata* (cross-hybrid between *M. indica* and *M. foetida*) (Teo et al. 2002) and *M. casturi* (Matra et al. 2021).

Close genetic relationship between *M. applanata* and *M. altissima has* been reported in a phylogenetic analysis conducted based on one chloroplast gene (*Maturase K*) while *M. indica* and *M. caloneura* have nested into two distinct clades (Hidayat et al. 2011). In our study, based on whole chloroplast genomes, *M. laijiwa*, *M. applanata*, *M. altissima* and *M. caloneura* were clustered with *M. indica* sharing 99.9% sequence similarity. These four wild relatives clustered with domesticated mango into a distinct clade even in concatenation-based nuclear phylogeny showing their close evolutionary relationship to *M. indica* whereas only *M. laijiwa* out of the above four species clustered separately in the coalescent approach. Furthermore, two of the *M. indica* cultivars (Kensington Pride and Tommy Atkins) were more closely related to *M. altissima* than *M. indica* cv. Alphonso failing to resolve *M. indica* from *M. altissima* based on nuclear gene sequences. A close evolutionary relationship between *M. altissima* and *M. indica* was also observed in the coalescence approach. Therefore, due to remarkably close evolutionary relationships observed among these species in chloroplast and nuclear phylogenies, we suggest these four wild relatives and cultivated mango are very closely related and might have shared or descended from the same common ancestor.

Considering domestication of *M. indica*, although a single domestication event has been reported based on historical records (Mukherjee 1972; Singh et al. 2016), two independent domestication events have also proposed for *M. indica* in India and Indochina (Bompard 2009). Based on a population genomics study, Warschefsky and von Wettberg (2019) suggested that mango domestication is a complex process and it may involve multiple domestication events and interspecific hybridization; two common phenomena observed in perennial fruit crop domestication. Their results also have indicated a high genetic diversity among *M. indica* cultivars distributed outside from the region where the mango was originated and a unique genetic diversity in Southeast Asian cultivars compared to other populations. Warschefsky and von Wettberg (2019) suggest that the origin and initial cultivation of mango may have taken place in Southeast Asia and further improvement and domestication may have occurred in India. In addition, cross-hybridization was highly likely to occur between wild relatives and *M. indica* at the early stages of domestication because of the presence of a high number of species and presence of evidence for crossbreeding. Thus, apart from descending from a common ancestor, cross-hybridization between *M. indica* and the four wild relatives is also a possible phenomenon that may have further contributed to the close evolutionary relationships observed in our study. However, this close evolutionary relationship could be further supported by including multiple replicates per species which is a limitation in this study.

*M. zeylanica*, is an endemic species to Sri Lanka which is only discovered in forests of wet and intermediate zones, showing scattered distribution. Close evolutionary relationship was observed in the concatenation-based nuclear phylogeny between *M. zeylanica* and *M. indica* in spite of having a distinct chloroplast genome when compared the variations between two species (with 44 INDELs, 122 SNPs and nine substitutions). Therefore, we hypothesized that cross hybridization might have occurred between an early lineage of *M. zeylanica* and *M. indica* or its close wild relative. Since the species have a distinct chloroplast genome, we assumed that during the cross, *M. indica* may have most likely acted as the pollen donor/ or paternal parent, resulting hybrids which carry the chloroplast genome *M. zeylanica* and nuclear genes of both *M. zeylanica* and *M. indica*/it’s close relative. The nuclear phylogeny/species tree based on the coalescence approach also showed a close relationship between *M. indica* and *M. zeylanica*. Clustering of *M. zeylanica* with *M. indica* in 14 individual gene trees suggested that *M. zeylanica* might have a hybrid origin. But as the BS/PP and LPP values are relatively low for this clade in individual gene trees as well as in both the consensus trees, it is also possible that the set of genes are not sufficiently variable to give a better resolution in the phylogeny. Therefore, it is difficult to make a conclusion about the hybrid origin for *M. zeylanica*.

Among six species (*M. casturi, M. laurina, M. pajang, M. odorata, M. percisiformis and M. hiemalis*) nested together in the chloroplast phylogeny, *M. casturi* is a cultivated species in Indonesia. This endemic species is only found in cultivation (Rhodes and Maxted 2016) and was proposed to be a natural hybrid between *M. indica* and *M. quadrifida* according to a SNP analysis (Warschefsky 2018). Since *M. casturi* has shown higher affinity to *M. indica* than to *M. quadrifida* instead of being direct intermediate between two species, it was further suggested that *M. casturi* is most likely a result of an F1 hybrid backcrossed with *M. indica* (Warschefsky 2018; (Warschefsky and von Wettberg 2023). Microsatellite marker based analysis showed broad genetic variation among four *M. casturi* accessions (Kasturi, Cuban, Pinari and Pelipisan) and DNA barcording based phylogenetic analysis suggested several species as ancesters for *M. casturi* (Matra et al. 2021). Genetic variation has also been confirmed between 16 accessions of *M. casturi* using SNP markers (N. Dillon, pers. comm.). Therefore a combination of microsatellite and DNA barcoading data support that *M. quadrifida* and *M. indica* was hybridised to result in M*. casturi* and F1 hybrids may have further hybridized with the ancestors of the parental species or multiple other *Mangifera* species to generate hybrid cultivars which have high genetic diversity (Matra et al. 2021). In our study, a close genetic relationship was observed between *M. casturi* and species in the domesticated clade of the concatenation-based nuclear phylogeny despite having distinct chloroplast genomes. In contrast, in the coalescence approach, *M.casturi* showed a relatively distant evolutionary relationship with *M. indica* both in species tree and in individual gene trees. Therefore, according to our results, coalescence-based nuclear phylogenies don’t strongly support the parentage of *M. indica* for *M. casturi*. Since a very low number of genes are shared between *M. indica* and *M. casturi*, if *M. indica* is one of the parents, it is not possible that *M. casturi* is a first-generation hybrid. Also, the absence of the data for other proposed parent (*M. quadrifida*) and other wild relatives limits analysing the input of *M. quadrifuda* and any other species for the hybrid origin of *M. casturi*.

*M. laurina* is a cultivated species in Indonesia where its wild distribution ranges from Myanmar, Cambodia, Vietnam and Malesia, Thailand to New Guinea. Analysis of *ITS* genomic region (Yonemori. et al. 2002) have revealed close evolutionary relationship between *M. laurina* and *M. indica.* Analysis of *Maturase K* chloroplast genomic region has differentiated Indonesia and Thailand specimens collected for *M. laurina*. Since common interspecific hybridization has been suggested for this species (Hartana 2010), it is possible that *M. laurina* may have cross hybridized with other species after intoroduction to the regions where it is widely cultivated. Due to the relatively close evolutionary relationship observed between *M. laurina* and *M. indica* in nuclear gene analysis despite the chloroplast genome being distinct, it might be possible to occur hybridization between early lineage of *M. laurina* and *M. indica*. Current data and results only support the close evolutionary relationship between the two species, but further analysis should be conducted with multiple samples for both species and also including more species in the genus.

Out of the remaining four species (*M. pajang, M. odorata, M. percisiformis and M. hiemalis*) clustered within the same main clade in chloroplast phylogeny, *M. pajang* is an endemic species originating from and cultivated in Borneo, Indonesia. *M. odorata* cultivated in Malaysia is a proposed hybrid between *M. indica* and *M. foetida* from which it showed more affinity to *M. foetida* than to *M. indica* (Teo et al. 2002; Yonemori. et al. 2002). Furthermore, closer genetic relationship of *M. foetida* to *M. pajang* compared to *M indica* has been suggested (Schnell and Knight Jr 1992). Having a distinct chloroplast genome and a distant phylogenetic relationship to *M. indica* in the chloroplast tree suggests that *M. odorata* might have captured chloroplast genome from *M. foetida* and ancestor of *M. indica* might have contributed as the male progenitor. In both concatenation and coalescence approaches for nuclear genes, *M. odorata* showed a relatively distant evolutionary relationship with *M. indica.* Since individual gene trees cluster the two species in four gene trees only with weak support, it is less likely that *M. odorata* is a first-generation hybrid. Though a small *M. indica* background was suggested for *M. odorata*, without being the other suggested parent, the hybrid status and parentage of *M. odorata* is inconclusive.

Another discrepancy observed from the chloroplast and nuclear trees is related to the position of *M. sylvatica*. Previous studies have revealed a close evolutionary relationship between *M. indica* and *M. sylvatica* based on restriction fragment length polymorphism (RFLP) (Eiadthong et al. 1999), *ITS* (Yonemori. et al. 2002) marker analysis and whole chloroplast genome analysis (Niu et al., 2021). In our study, although *M. sylvatica* showed a close genetic relationship with *M. indica* in chloroplast phylogeny, it nested with *M. hiemalis* in the nuclear phylogeny. *M. hiemalis* is an endemic species to China and *M. sylvatica* is also one of the cultivated species in China (Wang et al. 2020; Baul et al. 2016). Therefore, both species share the same geographical distribution. Since individual gene trees revealed clustering with *M. hiemalis* as sister taxa in 12 trees, *M. sylvatica* might have a hybrid origin where the hybridization occurred long ago, but the low BS values in individual gene trees does not strongly support this hypothesis.

Topological incongruence observed by chloroplast genome and single copy nuclear gene-based phylogenies reveal that there is a potential for inter-specific hybridization in the genus. But less BS values and weak resolution in gene trees of coalescence approach and low BS/PP/LPP support values in some of the branches of concatenation-based nuclear phylogeny and species tree are clear evidence that the nuclear genes are not well distinguished/ might not vary across the group of species studied. Less variability of nuclear genes and absence of one of the parents for proposed hybrids limited concluding about possible hybridization event/s occurred in the genus and hybrid of *M. odorata* and *M. casturi*. But even if we have hybrids in our dataset with the presence of both parents, phylogenies will show their close evolutionary relationships if it is a recent generation hybrid. Therefore, results of this study suggests that the whole group sufficiently closely related with each other so that we needed large amount of data to get a concatenated/consensus tree. The history of evolution of the species and hybridization is complex in the genus and requires more species to get better understanding. However, is it possible that out of 69 distinct species identified in the genus, some or many of them may have either domestication input or cross-hybridized with other wild relatives.

## 5. Conclusions

Our analysis of determining evolutionary relationships within the genus *Mangifera,* based on whole chloroplast genome and 47 single copy nuclear genes, revealed close genetic relationship among species and discrepancies between whole plastome and nuclear gene-based phylogenies. We suggest that the five species including *M. indica*, *M. altissima*, *M. applanata*, *M. caloneura*, *M. lalijiwa* are very closely related and might have descended from the same common ancestor due to their close evolutionary relationship. It was difficult to confirm the hybrid origin of *M. odorata* and *M. casturi* as suggested previously due to absence of one of the proposed parents within our dataset and due to clustering of the available proposed parent in only a low number of gene trees. Relatively high number of gene trees showed close evolutionary relationship between *M. zeylanica* and *M. indica,* and *M. sylvatica* and *M. hiemalis*. However, evidence did not strongly support the possible hybridization due to weak BS/PP and LPP supports in phylogenies. Moreover, it was observed that geographical proximity might have facilitated possible hybridization events. Despite limited number of species used in the study, it seems that evolution of species and hybridization in the genus *Mangifera* is a complex process. This is the first comparative analysis of evolutionary relationships within the genus with whole chloroplast genome and multiple nuclear genes. These findings provide an understanding about the nature of hybridization within the genus between wild and domesticated mango revealing potential domestication input into some species. Validation of hybridity and accuracy of evolutionary relationships within the genus can be highly supported and improved by adding more species including potential parents and sampling species from different geographical locations.

## 6. Authors contributions

**Robert J. Henry**: Conceptualization and design, Methodology, Supervision, Validation, Project administration, Funding acquisition, Resources and Writing - Review and Editing.

**Agnelo Furtado**: Conceptualization and design, Methodology, Investigation, Supervision, Validation, Formal analysis, Project administration, Resources, Software and Writing - Review and Editing.

**Natalie Dillon**: Conceptualization and design, Methodology, Supervision, Validation, Project administration, Funding acquisition, Resources, and Writing - Review and Editing.

**Ardashir Kharabian Masouleh**: Methodology, Supervision, Validation, Formal analysis, Resources, Software and Writing - Review and Editing

**WWM Upendra Kumari Wijesundara**: Formal analysis, Data curation, Investigation, Writing - original draft.

## 9. Data availability statement

All data supporting the findings of this study are available within the paper and its Supplementary Information.

## 10. Data Archiving Statement

Raw Illumina sequence read data were submitted to NCBI’ Short Read Archive (SRA) database under Bio project ID PRJNA940204 and under Bio sample ID’s SAMN33621737- SAMN33621749. Chloroplast genomes will be submitted to NCBI under Bio project ID PRJNA940204 and accession numbers will be provided.

## Acknowledgements and Funding

Authors acknowledge University of Queensland Research Computing Centre (UQ-RCC) for providing all the computation resources. David K Chandlee and Peter Salleras for providing leaf materials for the two species *M. pajang* and *M. caesia* respectively to use in the project. This project was funded by the Department of Agriculture and Fisheries-Queensland Alliance for Agriculture and Food Innovation (QAAFI) Collaboration Fund (Genomics of *Mangifera* species – HF11422) and the Hort Frontiers Advanced Production Systems Fund (National Tree Genomics – AS17000) as part of the Hort Frontiers strategic partnership initiative developed by Hort Innovation, with co-investment from the Queensland Government and contributions from the Australian Government.

## References

1. Acosta MC, Premoli AC (2010). Evidence of chloroplast capture in south American Nothofagus (subgenus *Nothofagus*, Nothofagaceae). Mol Phylogenet Evol 54(1): 235–242. 10.1016/j.ympev.2009.08.008.

2. Afinit MM (2017). Mrmckain/Fast-Plast: Fast-Plast V. 1.2. 6.

3. Ananda G, Norton S, Blomstedt C, Furtado A, Møller B, Gleadow R, Henry R (2021). Phylogenetic relationships in the *Sorghum* genus ba,sed on sequencing of the chloroplast and nuclear genes. The plant genome 14(3): e20123. 10.1002/tpg2.20123.

4. Ankenbrand MJ, Pfaff S, Terhoeven N, Qureischi M, Gündel M, Weiß CL, Hackl T, Förster F (2018). chloroExtractor: extraction and assembly of the chloroplast genome from whole genome shotgun data. J Open Source Softw 3(21):64. 10.21105/joss.00464.

5. Arumuganathan K, Earle E (1991). Nuclear DNA content of some important plant species. Plant Mol Biol Rep 9(3): 208–218. 10.1007/BF02672069.

6. Azim MK, Khan I. A, Zhang Y (2014). Characterization of mango (*Mangifera indica* L.) transcriptome and chloroplast genome. Plant Mol Biol 85(1-2): 193–208. 10.1007/s11103-014-0179-8.

7. Bakker FT, Lei D, Yu J, Mohammadin S, Wei Z, van de Kerke S, Gravendeel B, Nieuwenhuis M, Staats M, Alquezar-Planas DE (2016). Herbarium genomics: plastome sequence assembly from a range of herbarium specimens using an Iterative Organelle Genome Assembly pipeline. Biol J Linn Soc 117(1): 33–43. 10.1111/bij.12642.

8. Bally I, Akem C, Dillon N, Grice C, Lakhesar D, and Stockdale K (2010). Screening and breeding for genetic resistance to anthracnose in mango. IX International Mango Symposium 992: 239–244. 10.17660/ActaHortic.2013.992.31.

9. Bally IS, Bombarely A, Chambers AH, Cohen Y, Dillon NL, Innes DJ, Islas-Osuna MA, Kuhn DN, Mueller LA, Ophir R (2021). The ‘Tommy Atkins’ mango genome reveals candidate genes for fruit quality. BMC Plant Biol 21(1): 1–18. 10.1186/s12870-021-02858-1.

10. Bally IS, Dillon NL (2018). Mango (Mangifera indica L.) Breeding. In: Jameel M Al-Khayri, Shri Mohan Jain, Johnson DV (eds) Advances in Plant Breeding Strategies: Fruits. Springer, pp 811–896.. 10.1007/978-3-319-91944-7_20.

11. Bankevich A, Nurk S, Antipov D, Gurevich AA, Dvorkin M, Kulikov AS, Lesin VM, Nikolenko SI, Pham S, Prjibelski AD (2012). SPAdes: a new genome assembly algorithm and its applications to single-cell sequencing. J Comput Biol 19(5): 455–477. 10.1089/cmb.2012.0021.

12. Baul TK, Jahedul Alam M, Nath TK (2016). *Mangifera sylvatica* Roxb. in the forests of south-eastern Bangladesh: a potential underutilised tree for small-scale forestry. Small-Scale For 15(2): 149–158. 10.1007/s11842-015-9314-x.

13. Bompard J (1992). The genus *Mangifera* re-discovered: the potential contribution of wild species to mango cultivation. IV International Mango Symposium 341: 69–77. 10.17660/ActaHortic.1993.341.5.

14. Bompard J (2009). Taxonomy and systematics. The mango: Botany, production and uses. CAB International, Wallingford, 19-41. 10.1079/9781845934897.0019.

15. Charr J.-C, Garavito A, Guyeux C, Crouzillat D, Descombes P, Fournier C, Ly SN, Raharimalala EN, Rakotomalala JJ, Stoffelen P (2020). Complex evolutionary history of coffees revealed by full plastid genomes and 28,800 nuclear SNP analyses, with particular emphasis on *Coffea canephora* (Robusta coffee). Mol Phylogenet Evol 151: 106906. 10.1016/j.ympev.2020.106906.

16. Coissac E, Hollingsworth PM, Lavergne S, Taberlet P (2016). From barcodes to genomes: extending the concept of DNA barcoding. Mol Ecol 1423–1428. 10.1111/mec.13549.

17. Corriveau JL, Coleman AW (1988). Rapid screening method to detect potential biparental inheritance of plastid DNA and results for over 200 angiosperm species. Am J Bo 75(10): 1443–1458. 10.1002/j.1537-2197.1988.tb11219.x.

18. Darriba D, Taboada G, Doallo R, Posada D (2012). jModelTest 2: more models, new heuristics and parallel computing. Nat Methods 9(8): 772–772. 10.1038/nmeth.2109.

19. Degnan JH, Rosenberg NA (2009). Gene tree discordance, phylogenetic inference and the multispecies coalescent. Trends Ecol Evol 24(6): 332–340. 10.1016/j.tree.2009.01.009.

20. Dierckxsens N, Mardulyn P, Smits G (2017). NOVOPlasty: de novo assembly of organelle genomes from whole genome data. Nucleic Acids Res 45(4): e18–e18. 10.1093/nar/gkw955.

21. Dillon N, Innes D, Ming Y, Fang X, Li X, Gao X, Zhan RL, Hong Xia W, Bajaj P, Bally I, Kumar A (2016). Drafting the Kensington pride mango genome. In: Plant and Animal Genome Conference XXIV, San Diego, CA.

22. Dinesh M, Hemanth KV, Ravishankar K, Thangadurai D, Narayanaswamy P, Ali Q, Kambiranda D, Basha S (2011). Mangifera. In Wild crop relatives: genomic and breeding resources Springer, pp 61–74. 10.1007/978-3-642-20447-0_4.

23. Duarte JM, Wall PK, Edger PP, Landherr LL, Ma H, Pires PK, Leebens-Mack J, Depamphilis CW (2010). Identification of shared single-copy nuclear genes in *Arabidopsis*, *Populus*, *Vitis* and *Oryza* and their phylogenetic utility across various taxonomic levels. BMC Evol Biol 10(1): 1–18. 10.1186/1471-2148-10-61.

24. Eiadthong W, Yonemori K, Sugiura A, Utsunomiya N, Subhadrabandhu S (1999). Analysis of phylogenetic relationships in *Mangifera* by restriction site analysis of an amplified region of cpDNA. Sci Hortic 80(3- 4): 145–155. 10.1016/S0304-4238(98)00222-2.

25. FAOSTAT (2022). Food and Agriculture Organization of the United Nations. http://www.fao.org/faostat/en/#data/QC. Accessed 17 November 2022

26. Fitmawati F, Harahap SP, Sofiyanti N (2017). Phylogenetic analysis of mango (*Mangifera*) in Northern Sumatra based on gene sequences of cpDNA *trnL-F* intergenic spacer. Biodiversitas 18(2): 715–719. 10.13057/biodiv/d180239.

27. Fitmawati F, Hayati I, Sofiyanti N (2016). Using *ITS* as a molecular marker for Mangifera species identification in Central Sumatra. Biodiversitas 17(2): 653–656. 10.13057/biodiv/d170238.

28. Furtado A (2014). DNA extraction from vegetative tissue for next-generation sequencing. In Cereal genomics. Springer, pp 1–5. 10.1007/978-1-62703-715-0_1.

29. Guyeux C, Charr J-C, Tran HT, Furtado A, Henry RJ, Crouzillat D, Guyot R, Hamon P (2019). Evaluation of chloroplast genome annotation tools and application to analysis of the evolution of coffee species. PLoS One 14(6), e0216347. 10.1371/journal.pone.0216347.

30. Hartana A (2010). Phylogenetic study of *Mangifera laurina* and its related species using cpDNA *trnL-F* spacer markers. HAYATI J Biosci 17(1): 9–14. 10.4308/hjb.17.1.9.

31. Healey A, Lee D J, Furtado A, Henry RJ (2018). Evidence of inter-sectional chloroplast capture in Corymbia among sections *Torellianae* and *Maculatae*. Aust J Bot 66(5): 369–378. 10.1071/BT18028.

32. Hidayat T, Pancoro A, Kusumawaty D (2011). Utility of *matK* gene to assess evolutionary relationship of genus *Mangifera* (*Anacardiaceae*) in Indonesia and Thailand. Biotropia, 18(2): 74–80. 10.11598/btb.2011.18.2.41.

33. Hou D (1978). Florae Malesianae praecursores *LVI. Anacardiaceae*. Blumea: Biodiversity, Evolution and Biogeography of Plants, 24.

34. Iyer (1989). Recent advances in varietal improvement in mango. Acta Hortic 291: 109–132., 10.17660/ActaHortic.1991.291.14.

35. Jin J.-J, Yu W.-B, Yang J.-B, Song Y, DePamphilis C.W, Yi, T.-S, Li D.-Z (2020). GetOrganelle: a fast and versatile toolkit for accurate de novo assembly of organelle genomes. Genome Biol 21(1): 1–31. 10.1186/s13059-020-02154-5.

36. Jo S, Kim H.-W, Kim Y.-K, Sohn J.-Y, Cheon S.-H, Kim K.-J (2017). The complete plastome sequences of *Mangifera indica* L.(Anacardiaceae). Mitochondrial DNA B Resour 2(2): 698–700. 10.1080/23802359.2017.1390407.

37. Joly S, McLenachan PA, Lockhart PJ (2009). A statistical approach for distinguishing hybridization and incomplete lineage sorting. Am Nat 174(2): E54–E70. 10.1086/600082.

38. Katoh K, Standley DM (2013). MAFFT multiple sequence alignment software version 7: improvements in performance and usability. Mol Bio Evol 30(4): 772–780. 10.1093/molbev/mst010.

39. Kostermans AJGH, Bompard JM (1993). The mangoes : their botany, nomenclature, horticulture and utilization. Academic Press.

40. Letunic I, Bork P (2021). Interactive Tree Of Life (iTOL) v5: an online tool for phylogenetic tree display and annotation. Nucleic. Acids Res 49(W1), W293–W296. 10.1093/nar/gkab301.

41. Li W, Zhu X.-G, Zhang Q.-J, Li K, Zhang D, Shi C, Gao L-Z (2020). SMRT sequencing generates the chromosome-scale reference genome of tropical fruit mango, Mangifera indica. Biorxiv. 10.1101/2020.02.22.960880.

42. Li Z, De La Torre AR, Sterck L, Cánovas FM, Avila C, Merino I, Cabezas JA, Cervera MT, Ingvarsson PK, Van de Peer Y (2017). Single-copy genes as molecular markers for phylogenomic studies in seed plants. Genome Biol Evol 9(5): 1130–1147. 10.1093/gbe/evx070.

43. Liu B-B., Campbell CS, Hong D-Y, Wen J (2020). Phylogenetic relationships and chloroplast capture in the Amelanchier-Malacomeles-Peraphyllum clade (*Maleae*, *Rosaceae*): Evidence from chloroplast genome and nuclear ribosomal DNA data using genome skimming. Mol Phylogenet Evol 147: 106784. 10.1016/j.ympev.2020.106784.

44. Liu S, Wang X, Xie L, Tan M, Li Z, Su X, Zhang H, Misof B, Kjer KM, Tang M (2016). Mitochondrial capture enriches mito-DNA 100 fold, enabling PCR-free mitogenomics biodiversity analysis. Mol Ecol Resour 16(2): 470–479. 10.1111/1755-0998.12472.

45. Liu X, Wang Z, Shao W, Ye Z, Zhang J (2017). Phylogenetic and taxonomic status analyses of the Abaso section from multiple nuclear genes and plastid fragments reveal new insights into the North America origin of Populus (*Salicaceae*). Front Plant Sci 7: 2022. 10.3389/fpls.2016.02022

46. Luo R, Liu B, Xie Y, Li Z, Huang W, Yuan J, He G, Chen Y, Pan Q, Liu Y (2012). SOAPdenovo2: an empirically improved memory-efficient short-read de novo assembler. Gigascience 1(1): 2047-2217X-2041-2018. 10.1186/2047-217X-1-18.

47. Matra DD, Fathoni MAN, Majiidu M, Wicaksono H, Sriyono A, Gunawan G, Susanti H, Sari R, Fitmawati F, Siregar IZ (2021). The genetic variation and relationship among the natural hybrids of *Mangifera casturi* Kosterm. Sci Rep 11(1): 1–10. 10.1038/s41598-021-99381-y.

48. Moner AM, Furtado A, Henry RJ (2018). Chloroplast phylogeography of AA genome rice species. Mol Phylogenet Evol 127: 475–487. 10.1016/j.ympev.2018.05.002.

49. Moner AM, Furtado A, Henry RJ (2020). Two divergent chloroplast genome sequence clades captured in the domesticated rice gene pool may have significance for rice production. BMC Plant Biol 20(1): 1–9. 10.1186/s12870-020-02689-6.

50. Mukherjee S (1949b). A monograph on the genus *Mangifera*. Lloydia. 22: 73–136.

51. Mukherjee S (1972. Origin of mango (*Mangifera indica*). Econ Bot 26(3): 260–264. 10.1007/BF02861039.

52. Mukherjee S, Litz RE (2009). Introduction: botany and importance. In The mango: Botany, production and uses. pp 1–18. CAB International Wallingford UK. 10.1079/9781845934897.0001.

53. Mukherjee SK (1949a). The mango and its wild relatives. Sci Cult 26: 5-–9.

54. Niu Y, Gao C, Liu J (2021). Comparative analysis of the complete plastid genomes of *Mangifera* species and gene transfer between plastid and mitochondrial genomes. PeerJ 9: e10774. 10.7717/peerj.10774.

55. Niu Y, Gao C, Liu J (2022). Complete mitochondrial genomes of three *Mangifera* species, their genomic structure and gene transfer from chloroplast genomes. BMC Genom 23(1): 1–8. 10.1186/s12864-022-08383-1.

56. Odintsova MS, Yurina NP (2006). Chloroplast genomics of land plants and algae. In Biotechnological applications of photosynthetic proteins: biochips, biosensors and biodevices. Springer, pp 57–72. 10.1007/978-0-387-36672-2_6.

57. Rabah S, Lee C, Hajrah N, Makki R, Alharby H, Alhebshi AM, Sabir, J, Jansen R, Ruhlman T (2017). Plastome sequencing of ten nonmodel crop species uncovers a large insertion of mitochondrial DNA in cashew. The Plant Genome. 10(3): plantgenome2017-2003. 10.3835/plantgenome2017.03.0020.

58. Rebbeck CA, Leroi AM, Burt A (2011). Mitochondrial capture by a transmissible cancer. Science. 331(6015): 303–303. 10.1126/science.1197696.

59. Rhodes L, Maxted N (2016). Mangifera casturi. The IUCN Red List of Threatened Species. Accessed on 16 February 2023

60. Rieseberg LH (1995). The role of hybridization in evolution: old wine in new skins. Am J Bot 82(7): 944–953. 10.2307/2445981.

61. Rieseberg LH, Soltis D (1991). Phylogenetic consequences of cytoplasmic gene flow in plants. Evolutionary Trends in Plants 5(1): pp.65–84.

62. Ronquist F, Teslenko M, Van Der Mark P, Ayres DL, Darling A, Höhna S, Larget B, Liu L, Suchar MA, Huelsenbeck JP (2012). MrBayes 3.2: efficient Bayesian phylogenetic inference and model choice across a large model space. Syst Biol 61(3): 539–542. 10.1093/sysbio/sys029.

63. Saúco VG (2016). Mango rootstocks. Literature review and interviews.

64. Schnell R, Knight Jr R (1992). Genetic relationships among *Mangifera spp*. based on RAPD markers. Acta Hortic 341: 86–92 10.17660/ActaHortic.1993.341.7.

65. Singh N, Mahato A, Sharma N, Gaikwad K, Srivastava M, Tiwari K, Dogra V, Rawal S, Rajan S, Singh A (2014). A draft genome of the king of fruit, mango (*Mangifera indica* L.). Plant and Animal Genome XXII Conference. San Diego, CA, USA, pp 11–15

66. Singh NK, Mahato AK, Jayaswal PK, Singh A, Singh S, Singh N, Rai V, SV AM, Gaikwad K, Sharma N (2016). Origin, diversity and genome sequence of mango (*Mangifera indica* L.). Indian J Hist Sci 51(2.2): 355–368. 10.16943/IJHS%2F2016%2FV51I2.2%2F48449.

67. Singh NK, Mahato AK, Jayaswal PK, Singh S, Singh N, Yadav N, Rai, V, SV AM, Gaikwad K, Sharma N (2018). A reference genome assembly of the mango variety Amrapali (*Mangifera indica* L.). Plant and Animal Genome XXVI Conference San Diego, CA.

68. Smith RL, Sytsma KJ (1990). Evolution of *Populus nigra* (sect. Aigeiros): introgressive hybridization and the chloroplast contribution of *Populus alba* (sect. Populus). Am J Bo 77(9): 1176–1187. 10.2307/2444628.

69. Soorni A, Haak D, Zaitlin D, Bombarely A (2017). Organelle_PBA, a pipeline for assembling chloroplast and mitochondrial genomes from PacBio DNA sequencing data. BMC Genom 18(1): 1–8. 10.1186/s12864-016-3412-9.

70. Stamatakis A (2014). RAxML version 8: a tool for phylogenetic analysis and post-analysis of large phylogenies. Bioinformatics. 30(9): 1312–1313. 10.1093/bioinformatics/btu033.

71. Stegemann S, Keuthe M, Greiner S, Bock R (2012). Horizontal transfer of chloroplast genomes between plant species. Proc Natl Acad Sci USA 109(7): 2434–2438. 10.1073/pnas.1114076109.

72. Teo L, Kiew R, Set O, Lee S, Gan Y (2002). Hybrid status of kuwini, *Mangifera odorata* Griff.(*Anacardiaceae*) verified by amplified fragment length polymorphism. Mol Ecol 11(8): 1465–1469. 10.1046/j.1365-294x.2002.01550.x.

73. Tsitrone A, Kirkpatrick M, Levin DA (2003). A model for chloroplast capture. Evolution. 57(8): 1776–1782. 10.1111/j.0014-3820.2003.tb00585.x.

74. Tsutsui K, Suwa A, Sawada Ki, Kato T, Ohsawa TA, Watano Y (2009). Incongruence among mitochondrial, chloroplast and nuclear gene trees in Pinus subgenus *Strobus* (*Pinaceae*). J Plant Res 122(5): 509–521. 10.1007/s10265-009-0246-4.

75. Vasanthaiah HK, Ravishankar KV, Mukunda GK (2007). Mango. In C. Kole (ed) Fruits and Nuts. Springer, pp 303–323. 10.1007/978-3-540-34533-6_16.

76. Wang P, Luo Y, Huang J, Gao S, Zhu G, Dang Z, Gai J, Yang M, Zhu M, Zhang H (2020). The genome evolution and domestication of tropical fruit mango. Genome Biol 21(1): 1–17. 10.1186/s13059-020-01959-8.

77. Warschefsky E (2018). The evolution and domestication genetics of the mango genus, Mangifera(Anacardiaceae). FIU Electronic Theses and Dissertations. 3824. 10.25148/etd.FIDC006564.

78. Warschefsky EJ, von Wettberg EJ (2019). Population genomic analysis of mango (*Mangifera indica*) suggests a complex history of domestication. New Phytol 222(4): 2023–2037. 10.1111/nph.15731.

79. Warschefsky EJ, von Wettberg EJ (2023). The genetic composition of hybrid *Mangifera*. Biorxiv, 2023.2003. 2027.533847.

80. Wick RR, Schultz MB, Zobel J. Holt KE (2015). Bandage: interactive visualization of de novo genome assemblies. Bioinformatics. 31(20):, 3350-3352. 10.1093/bioinformatics/btv383.

81. Wolfe AD, Elisens WJ (1995). Evidence of chloroplast capture and pollen-mediated gene flow in Penstemon sect. Peltanthera (*Scrophulariaceae*). Syst Bot: 395–412. 10.1111/nph.15731.

82. Yi TS, Jin GH, Wen J (2015). Chloroplast capture and intra-and inter-continental biogeographic diversification in the Asian–New World disjunct plant genus Osmorhiza (*Apiaceae*). Mol Phylogenet Evol 85: 10–21. 10.1016/j.ympev.2014.09.028.

83. Yonemori K, Honsho C, Kanzaki S, Eiadthong W, Sugiura A (2002). Phylogenetic relationships of *Mangifera* species revealed by *ITS* sequences of nuclear ribosomal DNA and a possibility of their hybrid origin. Plant Syst. Evol. 231(1): 59–75. 10.1007/s006060200011.

84. Zhang C, Rabiee M, Sayyari, E, Mirarab S. (2018). ASTRAL-III: polynomial time species tree reconstruction from partially resolved gene trees. BMC bioinformatics 19(6): 15–30. 10.1186/s12859-018-2129-y.

85. Zhang N, Zeng L, Shan H, Ma H (2012). Highly conserved low-copy nuclear genes as effective markers for phylogenetic analyses in angiosperms. New Phytol 195(4): 923–937. 10.1111/j.1469-8137.2012.04212.x.

86. Zhang Y, Ou KW, Huang GD, Lu YF, Yang, GQ, Pang XH (2020). The complete chloroplast genome sequence of *Mangifera sylvatica* Roxb.(*Anacardiaceae*) and its phylogenetic analysis. Mitochondrial DNA B: Resour 5(1): 738–739. 10.1080/23802359.2020.1715286.

