## Supplementary material for "Determination of phylogenetic relationships in the genus *Mangifera* based on whole chloroplast genome and nuclear genome sequences": Figure S1

WWM Upendra Kumari Wijesundara (1), Agnelo Furtado (1), Natalie L. Dillon (2), Ardashir Kharabian Masouleh (1), Robert J Henry (1)
1 Queensland Alliance for Agriculture and Food Innovation, University of Queensland, Brisbane, 4072 Australia
2 Department of Agriculture and Fisheries, Mareeba, 4880, Australia

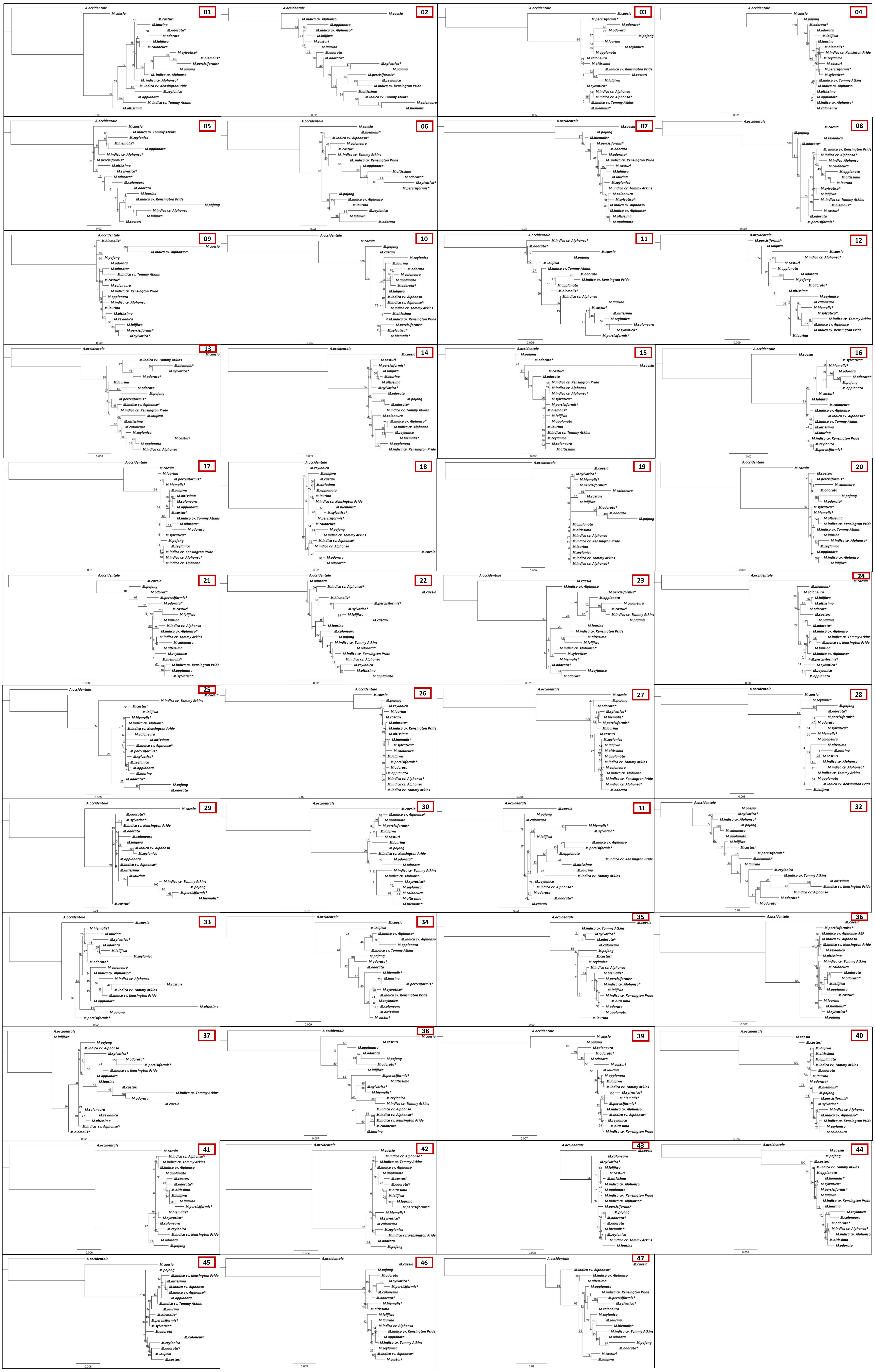


**Online Resource 5: Individual gene trees constructed for 47 single-copy nuclear genes.** Maximum Likelihood (ML) trees developed are shown here and the numbers associated with the branches of the trees are ML bootstrap values (/100).
